# Modular assembly of polysaccharide-degrading microbial communities in the ocean

**DOI:** 10.1101/387191

**Authors:** Tim N. Enke, Manoshi S. Datta, Julia Schwartzman, Nathan Cermak, Désirée Schmitz, Julien Barrere, Otto X. Cordero

## Abstract

Many complex biological systems such as metabolic networks can be divided into functional and organizational subunits, called modules, which provide the flexibility to assemble novel multi-functional hierarchies by a mix and match of simpler components. Here we show that polysaccharide-degrading microbial communities in the ocean can also assemble in a modular fashion. Using synthetic particles made of a variety of polysaccharides commonly found in the ocean, we showed that the particle colonization dynamics of natural bacterioplankton assemblages can be understood as the aggregation of species modules of two main types: a first module type made of narrow niche-range primary degraders, whose dynamics are controlled by particle polysaccharide composition, and a second module type containing broad niche-range, substrate-independent taxa whose dynamics are controlled by interspecific interactions, in particular cross-feeding via organic acids, amino acids and other metabolic byproducts. As a consequence of this modular logic, communities can be predicted to assemble by a sum of substrate-specific primary degrader modules, one for each complex polysaccharide in the particle, connected to a single broad-niche range consumer module. We validate this model by showing that a linear combination of the communities on single-polysaccharide particles accurately predicts community composition on mixed-polysaccharide particles. Our results suggest thus that the assembly of heterotrophic communities that degrade complex organic materials follow simple design principles that can be exploited to engineer heterotrophic microbiomes.

---

Many biological and technological systems are built by the integration of relatively autonomous parts, or modules, which can be rearranged to create larger multi-functional hierarchies^1,2^. Multi-domain proteins, regulatory networks or metabolic networks, to name a few, evolve in a modular fashion through the mix and match of simpler functional components, such as protein domains or metabolic pathways, that when combined form systems with more diverse functional repertoires^3^. In bacteria and *archaea*, for instance, new transcription factor proteins evolve by fusion of pre-existing signal-sensing protein domains, which monitor the intracellular environment, and DNA-binding protein domains, which modulate gene expression, enabling the rapid discovery of novel input-output pairs^4^. Likewise, bacteria and *archaea* can acquire new catabolic pathways via horizontal gene transfer from distant organisms and integrate them into a core network of metabolic reactions that generate energy and biomass precursors^5^. This ability to combine functional components or to plug them into existing infrastructures without disrupting their structure and function, is a key advantage of a modular design that enables the discovery of new functions by aggregating simpler components^6^.

Much like metabolic and gene networks, microbial communities might be conceptualized as interconnected systems, whereby populations of microbes interact via direct chemical communication, metabolic crossfeeding, etc. But while intracellular networks assemble via evolutionary processes, communities of microbes assemble and disassemble in ecological timescales via dispersal, colonization and growth^7^. Despite the frequent assembly and disassembly of microbial communities in variable environmental conditions, it is not well understood if communities can preserve a core structure across environments, modified only by the gain or loss of a few functional modules, or if instead, communities experience extensive species turnover due to a lack of modular organization. While the latter scenario has traditionally attracted strong interest as it can lead to alternative community states^8^, the potential for modular assembly in microbial communities has not been explored.

Addressing this important problem requires us to start by defining what we here will call an assembly module for an ecological community. By extension of the notion of an evolutionary module as used in the context of genome or metabolic network evolution^9^, we define an ecological assembly module as a group of taxa with similar dynamics and function, which can be integrated into various communities and perform a given metabolic process with minimal disturbance to the structure of the system. Although the term “module” has been applied in ecology to describe cohorts of species with dense patterns of interconnectivity in pairwise species interaction networks^10^, such a definition is independent of dynamics, and as such it need not relate to assembly modules as defined here, which in principle can be made of loosely connected species with similar function and dynamics.

In this study, we aim to establish whether the microbial communities that assemble on micro-scale particles of organic matter in the ocean do so in a modular fashion. In the ocean, much like in animal guts, heterotrophic microbes break down biopolymer particles, releasing and recycling bioavailable nutrients^11^. At the micro-scale, the decomposition of these complex carbohydrates depends on the assembly of communities on particles surfaces, which act as resources and community scaffolds^12,13^. Previous studies have shown that cross-feeding, in which an organism’s metabolic byproduct is the primary substrate of another organism, plays an important role in structuring communities on particles^14^. In this sense, particle-attached communities can be considered as self-organized metabolic collectives, where a number of species that co-colonize in an ordered fashion consume a primary resource, the particle biopolymer, and recycle byproducts through a series of trophic interactions.

To measure the potential for modular community assembly on particles, we performed controlled assembly experiments where communities are allowed to self-organize on particle surfaces, starting from the same species pool and in otherwise identical abiotic conditions, but changing the primary polysaccharide that makes the particle. This setup allows us to study how communities reorganize their structure as a function of perturbations in the initial substrate fed to the system, and to ask whether such reorganization reveals the presence of assembly modules. To implement these controlled community assembly experiments, we used model marine particles containing paramagnetic cores, ranging from 50 to 200 µm in diameter (Figure S1). Our particles were composed of one of four carbohydrates abundant in marine environments: chitin, alginate, agarose and carrageenan (Figure 1A), as well as combinations of these substrates. Chitin is frequently found in the shells of crustaceans such as copepods as well as on the membranes of diatoms^15,16^. Alginate is a structural component of the cell walls of brown algae, whereas agarose and carrageenan are enriched in seaweeds^17^. Particles of the different substrate types were incubated in natural seawater to perform community-capture experiments, where particles act as micro-scale community scaffolds that can be magnetically pulled down for genomic analysis or cultivation^13^.

**Figure 1.**
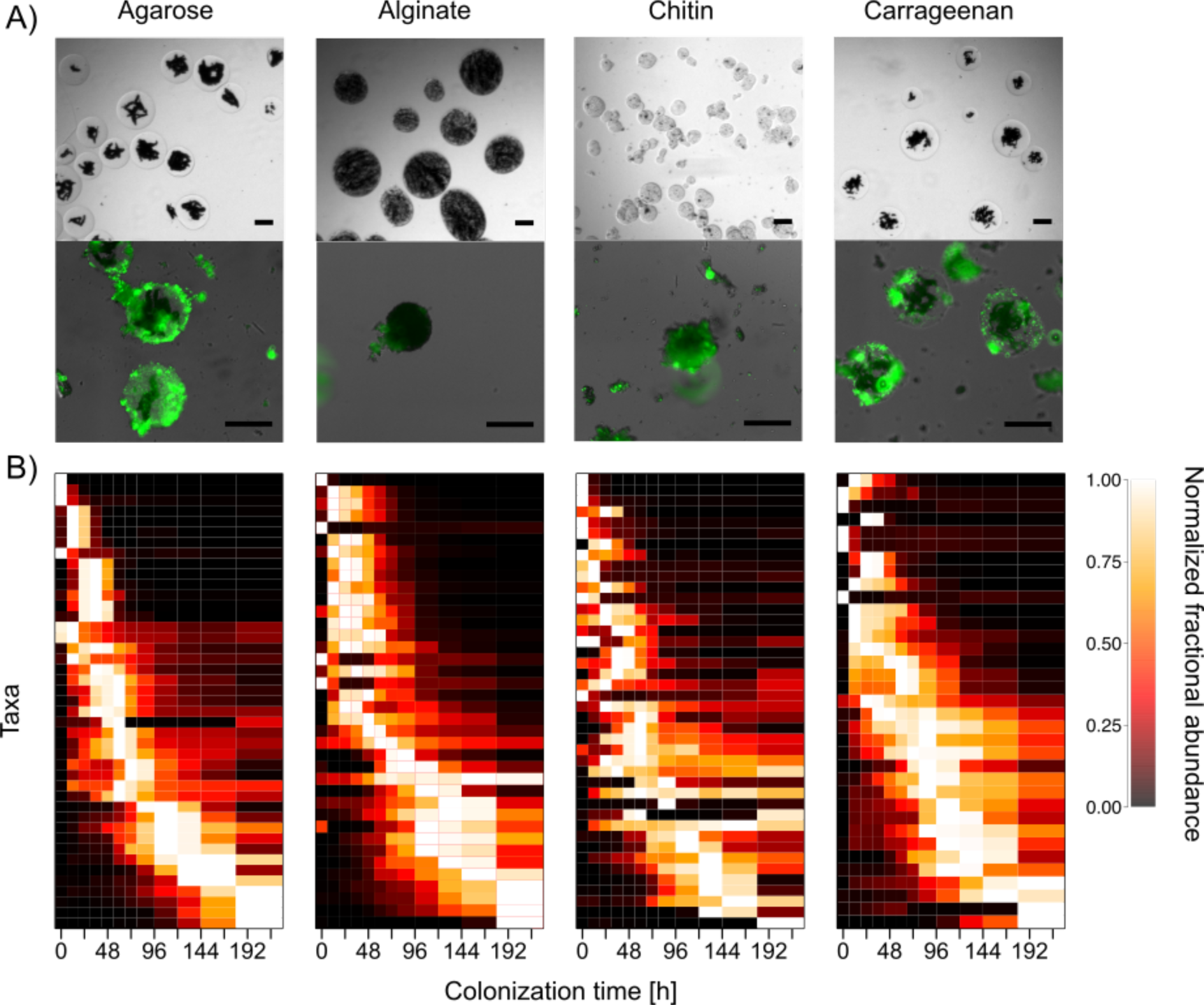
Rapid successional dynamics on four different marine polysaccharides. **A)**Paramagnetic hydrogel beads made of agarose, alginate, chitin, or carrageenan are incubated in natural, unfiltered coastal seawater. Upper panels are phase contrast images of the particles (with magnetite cores in black). Lower panels are fluorescence microscopy images of particles stained with Syto9 after 136 hours of incubation, revealing dense microbial communities on particle surfaces. Scale bar corresponds to 100 µm. **B)**Successional dynamics on each particle type. Taxa (rows) correspond to Amplicon Sequence Variants (ASVs) and are ordered by time at which they attain their maximum abundance. The data correspond to the relative frequencies of each taxon normalized by rows. Only ASVs whose maximum relative frequency is >1% are shown.

## Results

Previous work with chitin model particles has shown that community assembly proceeds in a reproducible succession, whereby early colonizers degrade chitin and facilitate the invasion of secondary consumers that lack enzymes required to hydrolyze chitin^14^. Across the four single-substrate particle types, we found that in all cases community assembly proceeded via rapid successional dynamics, indicating that the type assembly dynamics are not dependent on initial substrate. To characterize these dynamics, we collected ~1000 particles at each of twelve time points, from 0 to 204 hours, and sequenced their surface-attached communities using 16S rRNA gene amplicon sequencing (Methods). With this data we calculated Amplicon sequence Variants(ASVs) using the DADA2 pipeline^18^, identified the most abundant ASVs – comprising at least 1% of sequenced reads for at least one time point – and ordered them by the time at which they reached their maximum abundance within the communities (Figure 1B). On all four particle types tested, most of taxa present at high abundance in the first 12 hours decline substantially in abundance by 72-96 hours, indicating a remarkably similar rapid community turnover.

Despite the overall similarity in colonization dynamics across particle types, the abundance and dynamics of individual ASVs on different particle types was not necessarily conserved. To quantify differences in ASV abundance across particle types, we calculated a “niche breadth” index for the ASVs. To this end, for each ASV, *i*, and each particle type, *j*, we computed the geometric mean frequency over time, *f*_*ij*_, renormalized the mean frequencies so ∑_*j*_*f*_*ij*_ = 1 and calculated the entropy of the mean ASV abundance over particle types, −∑_*j*_*f*_*ij*_log_2_(*f*_*ij*_)(Methods). The entropy represents an index that described how uniformly the ASV was distributed over the four substrates. ASVs that appeared only on one particle type had a niche breadth score = 0,whereas ASVs that were equally prevalent across all particle types (*f_ij_* = 0.25 ∀*j*) had a niche breadth score index of 2.

We found that within particle-associated communities the distribution of the niche breadth indexes was bimodal (top histogram in Figure 2A). Using a Gaussian mixture model to cluster ASVs by distribution mode (Methods), we found that 36% of the ASVs grouped into a cluster of narrow-range taxa (niche breadth score < 0.18) and 42% into a cluster of broad-range taxa (niche breadth score > 1.52). Moreover, an unsupervised hierarchical clustering of ASVs based on their temporal dynamics across particle types allowed us to further partition narrow-range taxa by the substrate they appeared on (heat map in Figure 2A). The best partitioning of the data divides ASVs into five natural blocks, one for the broad-range taxon set and one block of narrow-range taxa for each of the four particle types (Methods). The broad-range block encompassed organisms that were not only highly prevalent across all particle types, but whose dynamics were highly correlated across substrates (average Spearman correlation = 0.54 across four particle types, Figures S2-S3). On average, these broad-range taxa increased in frequency towards later time points, causing community composition across particle types to first diverge due to the colonization of narrow-range species (reaching maximum divergence at ~24h) before converging to a set of broad-range taxa (Figure S4, S5). Overall, the comparison of the assembly dynamics across particle types shows that community assembly on particles can be coarse-grained in terms of blocks of species with correlated dynamics, representing putative assembly modules, which are either highly specific or unspecific to the primary polysaccharide that feeds the community.

**Figure 2.**
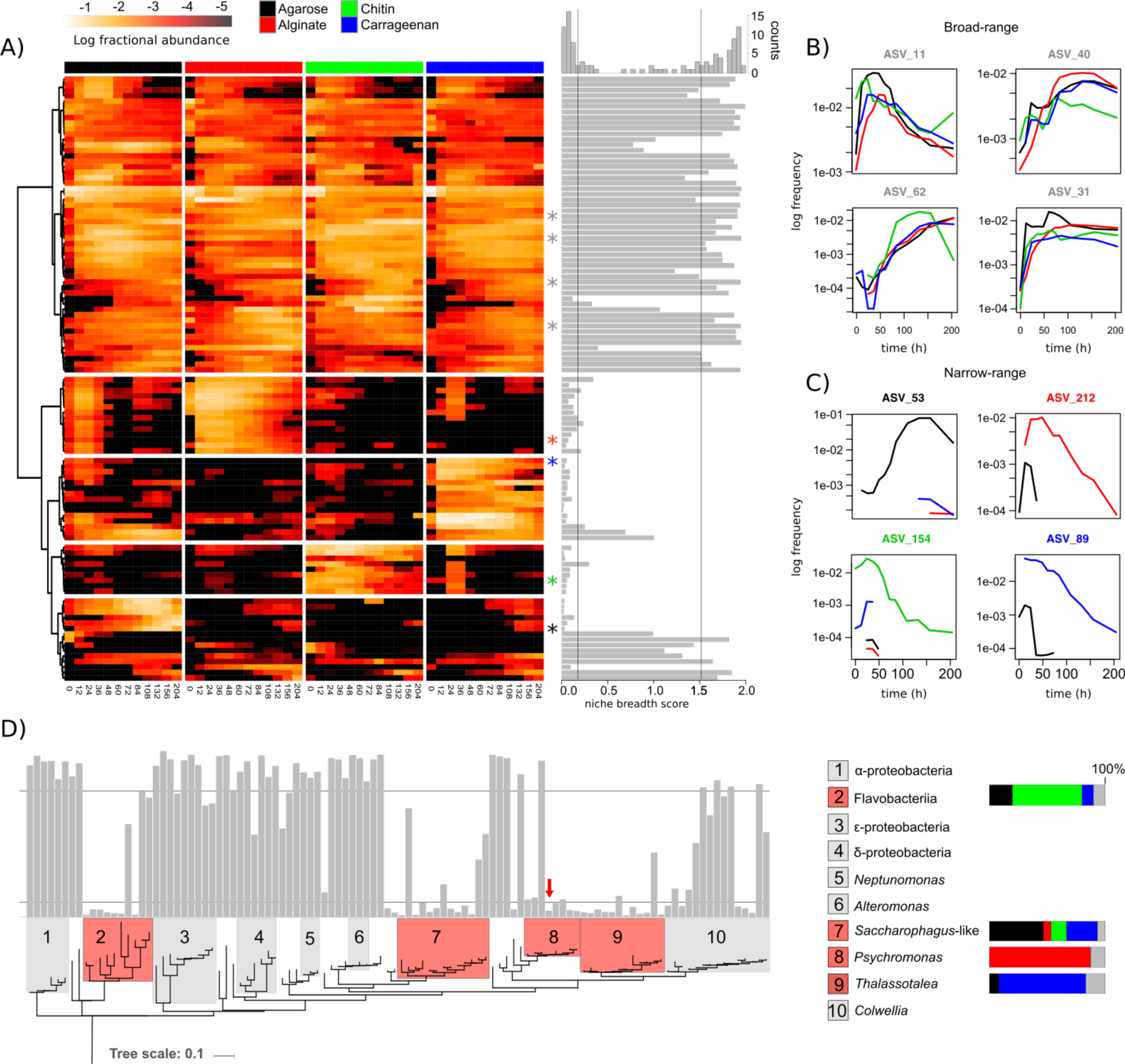
Unsupervised detection of narrow-range and broad-range taxa. **A**. Clustering of taxa by occurrence across four particle types. The data show that taxa can be divided in a large cluster of broad-range taxa and smaller clusters of narrow-range taxa. The niche breadth score (gray bars) shows that the distribution of niche breadths is bimodal (see histogram on top). **B-C)** Dynamics of broad-range ASVs (B), four cases, and narrow-range ASVs (C), one case for each substrate. The position of these specific ASVs are marked with an asterisk (*) in the heat map in A. The color of the asterisk corresponds to the color coding of substrates (legend in panel A), with the broad-range ASVs colored in gray. Figures S2 and S3 show the dynamics of all broad-range and narrow-range taxa, respectively. **D)** Phylogenetic distribution of narrow- and broad-range taxa. Phylogenetic clusters marked with numbers correspond to the largest monophyletic clades, defined at the class level for groups 1,2 and 3, and at the genus level for groups 4-10, all of which fall within the γ-proteobacteria class. In red are those monophyletic clades with a high incidence of narrow-range taxa (>50%). Tree rooted with Sulfolobus as outgroup (not shown). Red arrow points to the position of psychB3M02, the alginate degrader mentioned in the main text. Panel on the right shows the taxonomic description of the clades and the distribution of ASVs across each of the four narrow-range clusters with colored bar plots.

A phylogeny of the ASVs showed that narrow- and broad-range blocks were associated with distinct taxonomic groups and distinct metabolic potentials. Narrow-range blocks mapped primarily to four taxonomic groups: the family *Flavobacteriaceae*, which contributed to most chitin-associated ASVs, the genera *Sacharophagus* and its close relatives (e.g. *Terednibacter*), contributing most carrageenan-associated ASVs, *Psychromonas*, with virtually all ASVs in the alginate block, and *Thalassotalea*, contributing most carrageenan-associated ASVs (Figure 2D, clades 2,7,8 and 9). Marine bacteria of the class *Flavobacteriia* and the genus *Sacharophagus* are among the most well-known degraders of polysaccharides in the ocean^19–22^, suggesting that these narrow-range taxa are specialized primary degraders. To gain further insight into the genomic and metabolic differences between narrow and broad-range taxa that could explain their dynamics, we cultured 874 bacterial isolates from particles and sequenced their 16S rRNA V4 region (Methods and SOM). Out of these, 247 isolates had a 100% identity match to 12 broad-range ASVs. Only 2, however, mapped to 2 narrow-range ASVs (SOM). We focused our efforts on one of these narrow-range isolates, which we named psychB3M02, and belonged to the genus *Psychromonas* in the alginate-specific block (marked with a red arrow in Figure 2D). In agreement with its specific association with alginate particles, psychB3M02 was able to grow on alginate as sole carbon source (Figure 3A). Moreover, HMM-based searches of glycosyl hydrolase (GH) and polysaccharide lyase (PL) families against its genome identified multiple copies of alginate lyases (PL7, 8 copies), and oligoalginate lyases (PL15, PL17, 4 copies), but found no other genes coding secreted enzymes for degrading other marine polysaccharides such as chitin (GH18, GH19, GH20) or agarose (GH16) (Table S1). The absence of other polysaccharide degrading enzymes suggests that psychB3M02 has a specialized role as a primary degrader of alginate, in agreement with its narrow niche range.

**Figure 3.**
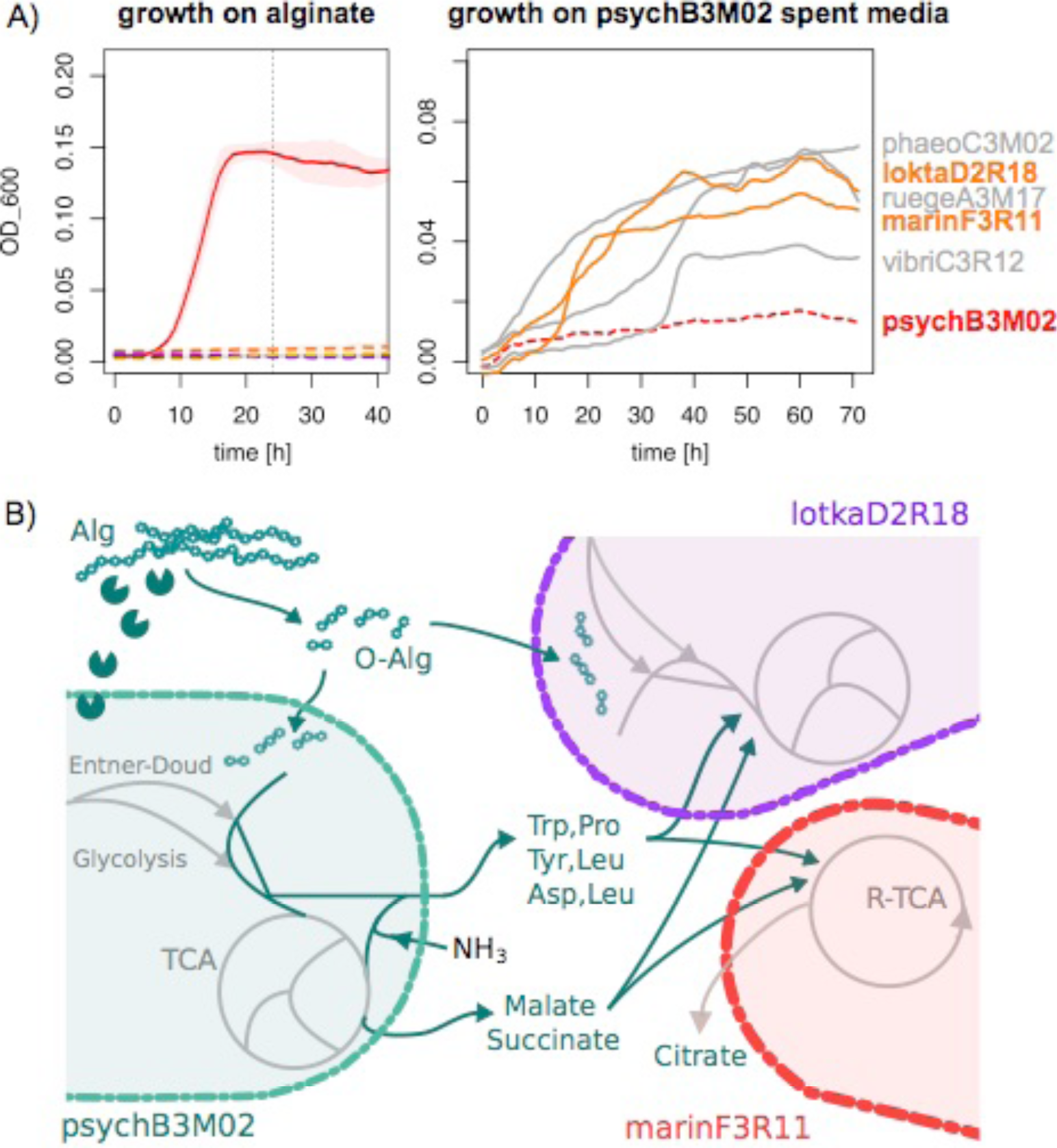
Facilitation of the broad-range module is generic and mediated by multiple amino acid and organic acid excretions. **A)**. Growth curves of a narrow-range degrader, psychB3M02, and 5 broad-range non-degraders on alginate (left) and on spent media of psychB3M02. **B)**Model of possible cross-feeding pathways inferred from full genomes of psychB3M02, lotkaD2R18 and marinF3R11, as well as from targeted metabolomics data (Tables S2-S4).

By contrast, none of three isolates of the *Rhodobacteraceae* (α-proteobacteria), a clade exclusively found in the broad-range block (Figure 2D, clade 1), encoded genes to produce hydrolytic enzymes (Table S1). Two members of this clade, however, a *Loktanella*, lotkaD2R18, and a *Ruegeria*, ruegeA3M17, had the machinery to import and utilize oligosaccharides of alginate and chitin, respectively, suggesting a potential role as ‘free-riders’. By contrast, the third organism, phaeoC3M10, classified as a *Phaeobacter*, had no genes to convert cytoplasmic intermediates into central metabolic substrates, indicating that this strain cannot harvest oligosaccharides and instead relies on metabolic intermediates released by other members of the community. To experimentally assess the potential for facilitation between narrow- and broad-range taxa, we collected spent media from psychB3M02 grown to peak cell density on alginate as the sole carbon source and asked whether this media would support growth of a panel of five broad range taxa that were unable to degrade and grow on alginate by themselves. We tested the three *Rhodobacteraceae* discussed above plus a *Marinobacter* and a *Vibrio* (Fig. 3A). In accordance with our expectation, all five broad-range taxa were able to grow on the spent media, even without supplementing it with additional nutrients (Figure 3A). This confirms that in an environment where alginate is the sole carbon source, narrow-range alginate degraders can facilitate the growth of broad-range, non-degrading taxa.

To learn more about the exact mechanisms of facilitation and its apparent non-specific nature, we performed a targeted metabolomic analysis^23^of psychB3M02’s spent media before and after growth of non-degrading broad range taxa (Methods), which showed that non-degraders support their growth by taking up multiple small metabolic byproducts. For this analysis, we picked two non-degrading strains whose genomes suggested divergent metabolic capabilities: the *Loktanella* lotkaD2R18 and the *Marinobacter* marinF3R11. We identified compounds that were produced by psychB3M02 and consumed by one of the non-degraders in at least two out of three replicates. Out of 82 possible compounds, we detected 11 compounds that fulfilled this criterion: these included six amino acids (Figure 3B), the amino acid precursor 3-methyl-2-oxopentanoic acid, TCA cycle intermediates malate and succinate, nucleosides and nucleotides (Tables S2-S4). This general consumption of multiple metabolic intermediates was observed for both marinF3R11 and lotkaD2R18. Some metabolites that could support growth of non-degraders were also released to the medium by non-degraders (Figure 3B). In particular, marinF3R11 secreted citrate, consistent with the prediction that this organism uses a reductive TCA cycle (Table S1). Overall, these data suggest that simultaneous utilization of a variety of metabolic intermediates is a robust ecological strategy for broad-range organisms, which could enable their growth in a manner that is not specific to the carbohydrate fed to the community.

Having identified five distinct functional components, one for each primary substrate and one for the group of cross-feeding broad-range taxa, as well as their mechanism of interaction, we asked whether communities capable of degrading multiple polysaccharides could be assembled in modular fashion, that is, by a simple aggregation of polysaccharide-specific modules. If this were the case, we would expect that the composition and dynamics of a community of higher complexity, capable of degrading multiple primary substrates, should be well approximated by a simple linear combination of the components we have identified. To test this hypothesis, we examined community assembly dynamics on particles made of substrate mixtures and compared these dynamics to the one observed on the corresponding single substrate particles. In particular, we tested two mixed particle types: agarose-alginate and agarose-carrageenan (50% of each substrate by mass), which were incubated in the same seawater and conditions used for single substrate particles.

Consistent with the notion of community assembly by aggregation of polysaccharide-specific modules, a simple linear combination of the species abundances on each single substrate accurately predicted the composition of communities assembled on mixed particles (Figure 4A). To quantify this, we fitted the vector of ASV geometric mean frequencies on the mixed particles with a linear combination of the vectors of the corresponding single substrate particles (SOM). The best fitting linear model for the agarose-alginate mixture, 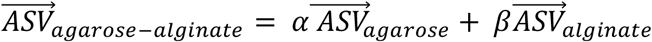 had an R^2^ of 0.84, and the corresponding model for agarose-carrageenan an R^2^of 0.74, showing that a linear combination had high-predictive power (Figure 4A). To rule out the possibility that the result was driven by broad-range taxa, we calculated the Spearman correlation coefficient between model and data only for the relevant narrow-range ASVs, finding values of 0.75 and 0.83 for the agarose-alginate and agarose-carrageenan communities, respectively, showing that the results hold for narrow-range taxa alone. Furthermore, we also fitted a model with an explicit interaction term to test if this would improve the results. We found that for the agarose-alginate such a nonlinear model had an inferior goodness-of-fit compared to the simple linear combination (SOM). In the case of agarose-carrageenan particles, the nonlinear model (nlm) improves the fit relative to the linear model (lm), but only marginally (R^2^ = 0.76 vs 0.74 in the lm) and the model is only weakly nonlinear (nlm~ lm^0.98^) (SOM). This analysis was based only on the average abundance of the ASV over time, however, when we considered their dynamics we found these were also highly correlated between single and mixed particles, in a manner consistent with a model of community assembly by simple linear aggregation of ecological modules (Figure 4BC). Across all alginate, agarose and carrageenan narrow range ASVs, the median Spearman correlation between the single-and mixed-substrate time dynamics ranges between 0.65 and 0.96 (Table S5). Overall, these results show that there is minimal interference between narrow-range modules, such that a linear combination model provides a good prediction of the assembly of the community on mixed substrate particles. This lack of interference, combined with the ability of broad-range modules to “plug-in” to narrow-range ones in a substrate independent manner, leads to the modular assembly of polysaccharide-degrading communities with a larger repertoire of metabolic functions (Figure 4D)

**Figure 4.**
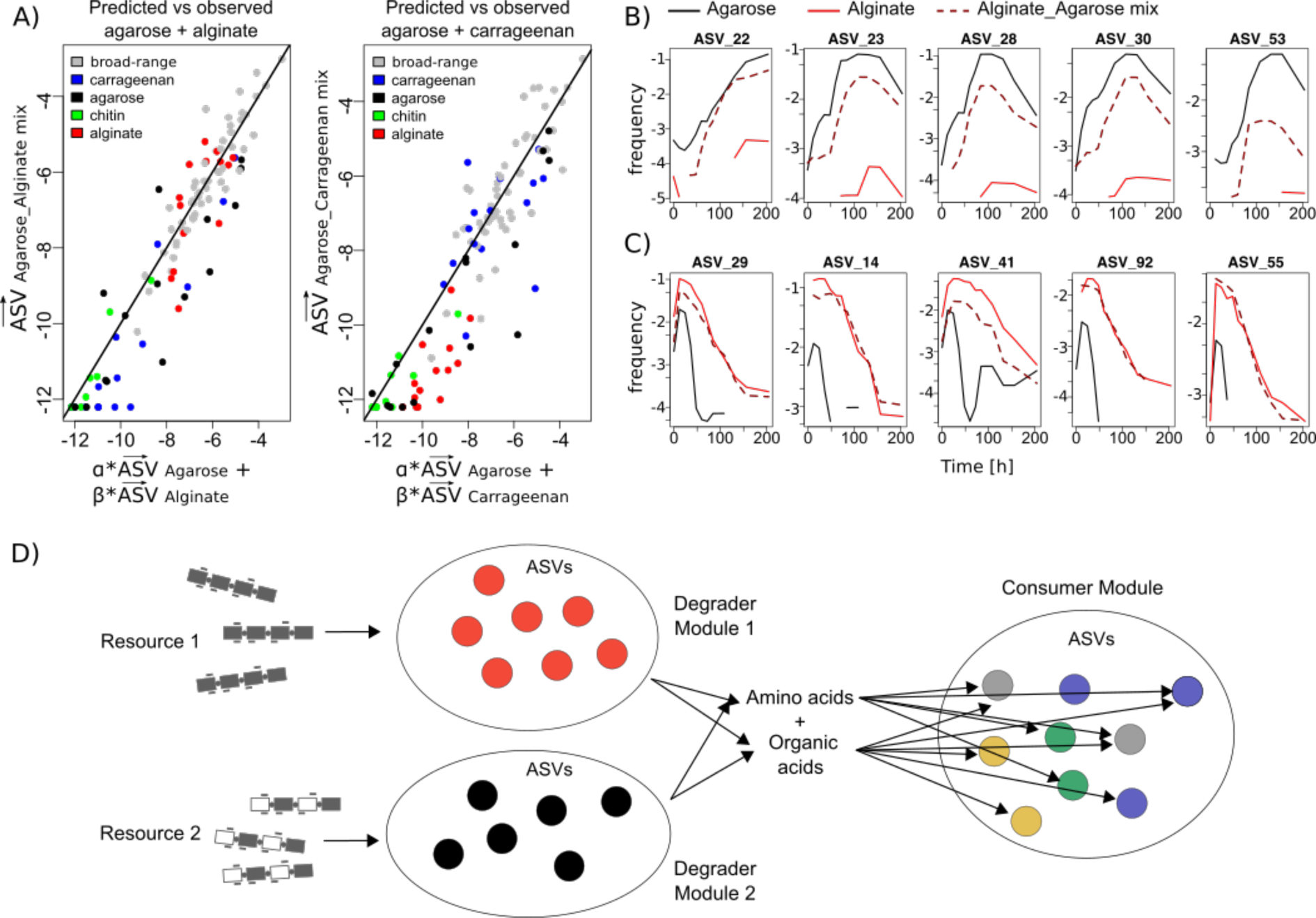
Communities assemble by linear combination of modules. **A)** ASV frequencies in mixed particles plotted in log-log scale against the predicted ASV frequencies, based on a linear combination of single substrate vectors. The fitted coefficients are α = 0.67, β = 0.40 for agarose-alginate and α = 0.89, β = 0.11 for agarose-carrageenan. **B-C)**. Similar ASV trajectories in mixed vs. single substrate particles for agarose (B) and alginate (C) specific ASVs. Solid lines depict trajectories in single substrate particles and dashed lines in mixed particles. The median Spearman correlation between the dynamics of agarose-specific ASVs on single and mixed substrate particles is 0.86 (B), and for the alginate-specific ASVs 0.96 (C) (Table S5). **D**) Model of modular assembly, which mirrors the structure of metabolic pathways. Peripheral, narrow-range modules perform the degradation of complex biopolymers, whereas the core, broad-range module processes simple metabolic intermediates.

In this study, we have shown that, despite the myriad species present in polysaccharide degrading communities, these systems can be coarse-grained into functional components, which assemble modularly into a variety of arrangements giving rise to communities of different functional complexity. Modules are divided into two classes, those encompassing species capable of breaking down polymers and those that encompass species that can live off metabolic byproducts. This subdivision mirrors the modular organization of metabolic pathways, in which sets of genes coding for hydrolytic enzymes, transporters, etc. can be horizontally acquired by an organism and integrated into its metabolic network as long as the products of the metabolic conversions performed by the integrated module are compatible with core metabolic pathways, such as glycolysis^5^. In this way, simple metabolic byproducts act as a common interface for pathways to interact, enabling organisms to acquire a variety of degradation modules, and to quickly modify their resource utilization profile^5,24^. Similarly, our data suggests that ecological modules of particle degrading bacteria interact with modules of byproduct consumers through multiple central metabolites, which form a common interface that might allow consumers to grow regardless of the initial polysaccharide fed to the community (Figure 4D). Interestingly, modules can have characteristic phylogenetic distributions, with taxa such the genus *Psychromonas* or the family *Flavobacteriaceae* being strongly associated with specific substrates. However, these associations between taxonomy and function need not necessarily be stable, as members of these taxonomic groups have been found that are specialized to degrade different polysaccharides^25^. In sum, our work suggests that modularity could play an important role in the assembly of natural microbial communities, and that it is a property that can emerge from the underlying metabolic organization of the community members. Future work should seek to validate this principle across more functional dimensions and to explore its applicability in the design of synthetic consortia.

## Methods

### Sampling and incubation

Coastal ocean surface water samples were collected in 2015 from Canoe Beach, Nahant, Massachusetts, USA; 42° 25’11.5’’N, 70° 54’26.0’’W. For each particle type, we set up triplicate 800 ml seawater incubations with model particles, using 1L wide-mouth Nalgene bottles. Particles, which had been stored in artificial sea water (Sigma, #S9883) with 20 % ethanol, were washed twice with artificial seawater to remove the ethanol and inoculated at a concentration of 100 particles per mL. Bottles were rotated overhead at room temperature and a speed of 7.5 rpm for 10 days. At t = 0, 12, 24, 36, 48, 60, 72, 108, 132, 156, 180, 204 hours, 10 mL (~1000 particles) were sampled from each replicate incubation and particles collected by magnetic separation for DNA sequencing and isolation.

### 16S amplicon data analysis

16SrRNA sequencing libraries were prepared in house according as in ^14^to the protocol described in the SOM. Sequencing was done at the BioMicroCenter at MIT. To identify Amplicon Sequence Variants (ASVs) from the 16SrRNA amplicon reads, we used the DADA2 pipeline^26^. We developed a pipeline based on the DADA2 developers’ ["Big Data: Paired-end" workflow] (http://benjjneb.github.io/dada2/bigdata_paired.html), which has been deposited in a public repository on [Github](https://github.mit.edu/josephe/dada2_pipeline). Briefly, a parametric error model is learned from the sequencing data, using a subset of two million reads drawn randomly from all those sequenced. Then, this error model is used to "denoise" samples by identifying erroneous sequence variants and combining them with the sequence variant from which they most likely originated. All other read processing steps -- including merging paired-end reads, trimming primer sequences, and dereplicating reads -- were performed with functions from the R Bioconductor "dada2" package.

For our analysis, we focused on the abundant ASVs, defined as those with a frequency > 1% in at least one sample across all samples, including replicates, time points and single substrate particle types. The resulting 107 ASVs were used throughout our analysis. Replicates were combined by calculating the weighted average frequency for every ASV, using the read counts of that sample as weights. We smoothed the data with a running median filter, window size = 3 and renormalized to work with mean frequencies.

### Niche breadth index

To study the prevalence of each ASV across different particle types, we devise a niche breadth index. We calculated the geometric mean frequency ASV on a particle type, *f_ij_ = e*^*<Iog(fij(t))>*^, where *f_ij_(t)* is the frequency of ASV *i* at time *t* on particle type *j*. We added pseudo counts (10^-6^) to *f_ij_(t)* to account for zeroes. With the normalized geometric mean frequencies, 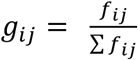 we calculated a niche breadth index over *j* using the entropy: − Σ*g*_*i*_log (*g*_*i*_). We use the R function Mclust to group our ASVs into three optimal groups according to their niche breadth index. The niche breadth index cutoff values for the groups are < 0.18 and > 1.52. The three resulting groups have 38, 24 and 45 members, respectively.

### Hierarchical clustering of ASV trajectories

We cluster the most abundant ASVs based on their log-transformed frequencies across all time-points and all particle types. We used the R function *hclust* with the clustering method ’*ward.D’* and Euclidean distances. To evaluate the best cutoff for our hierarchical clustering, we cut the tree generated by ’hclust’ into 2-15 groups using the ’cutree’ function in R. We used use the silhouette function from the R package ‘cluster’ to evaluate the clusters generated. Our analysis shows that 5 clusters are the optimal partitioning of our data.

### Phylogenetic tree of ASVs

To create a phylogenetic tree of the top 1% ASVs, we first aligned the 16S V4V5 sequences on Silva’s SINA alignment server (http://www.arb-silva.de/aligner/) with standard settings, the option *Search and Classify* enabled with *minimum identity with query sequence* = 0.9 and *classification: rdp*. After removing non-informative positions from the alignment we used FastTree 2.1^27^with the options -gtr -n) to infer an approximate maximum-likelihood phylogenetic tree which we upload to iTol^28^.

### Isolation of bacteria attached to particles

After 1.5, 3.5 and 6.5 days of incubation, particles were sampled, separated from the sea water and washed as described above and split into 1:1, 1:10 and 1:100 dilutions in artificial sea water (Sigma, #S9883). Dilutions were vortexed for 20 seconds and plated using glass beads (Zymo #S1001) on 1.5 % agar (BD #214010) plates with (1) Marine Broth 2216 (Difco #279110) or (2) Tibbles-Rawling minimal media as described in ^14^with carbon sources specific for the particle type: 0.05 % alginate, 0.04 % carrageenan, 0.1 % glucosamine, or plain agar. Following two days of incubation at room temperature, at least 16 colonies per particle and plate carbon source type were picked and re-streaked twice on Marine Broth 2216 1.5 % agar plates for purification. To obtain stocks, purified isolates were grown in deep well plates with liquid Marine Broth 2216 for 48 hours, shaking at 300 rpm at room temperature. The liquid culture was frozen at -80 °C for further characterization. Taxonomic classification was done using the 16S rRNA and the RDP database (https://rdp.cme.msu.edu/classifier/classifier.jsp)^29^.

### Crossfeeding experiments

The alginate-degrading strain psychB3M02 was streaked on Marine Broth 2216 1.5% agar plates and incubated at 25°C. After 48 hours single colonies were picked and grown in liquid Marine Broth at 25°C. After 48 hours, cells were pelleted and washed with Tibbles-Rawling minimal media twice. PsychB3M02 cells were then transferred at a starting OD of 0.005 to Tibbles-Rawling minimal media with 0.15% alginate (Sigma, #A1112) as the sole carbon source, and incubated in 10 mL volumes at 20°C and with overhead rotation. After 24h, the spent media was harvested by gently pelleting the cells (3000 rcf for 10 min) and filtering the supernatant through a 0.2 µm syringe filter. The five alginate non-degraders were pre-grown and harvested in a similar manner and transferred to fresh raw spent media at a starting OD of 0.005 in 200µl volumes. Growth was measured using OD600 on a Synergy2 microplate reader (BioTek).

### Genome sequencing

For selected isolates from our collection, genomic DNA was extracted from a liquid overnight culture in Marine Broth 2216 (Difco #279110) using the Agincourt DNA Advance Kit (Beckman Coulter #A48705). Genomes were sequenced using the Nextera DNA Library Preparation Kit (Illumina #FC-121-1031)^30^. Sequencing was performed on an Illumina HiSeq 2500 (250x250 bp paired-end reads) at the Whitehead Institute for Biomedical Research (MIT, Cambridge, MA, U.S.A.). Genomes were assembled using CLC Genomics Workbench 11 (Qiagen), curated using CheckM^31^. Open reading frames were annotated using the RAST pipeline^32^and the CAZY database^33^, run from the dbCAN2 server^34^. Sequences were deposited in project PRJNA478695.

### Metabolomics

Metabolomics was performed at the Microbial Biogeochemistry Group at the Woods Hole Oceanographic Institution. To extract the metabolites from the spent media, the filtrate was acidified to a pH ~3 using 12 M hydrochloric acid and the extracellular organic compounds extracted using Bond Elut PPL cartridges (1 g/6 ml sized cartridges, Agilent) following the protocol of Dittmar et al.^35^as modified by Longnecker^36^. Dissolved organic matter was eluted from the cartridges using 100% methanol. The resulting organic matter extracts were analyzed using targeted mass spectrometry. Briefly, the extracts for targeted analysis were re-dissolved in 95:5 (v/v) water:acetonitrile with deuterated biotin (final concentration 0.05 mg ml^-1^). Samples were then analyzed by ultra performance liquid chromatography (Accela Open Autosampler and Accela 1250 Pump, Thermo Scientific) coupled to a heated electrospray ionization source (H-ESI) and a triple quadrupole mass spectrometer (TSQ Vantage, Thermo Scientific) operated under selected reaction monitoring (SRM) mode. Chromatographic separation was performed on a Waters Acquity HSS T3 column (2.1 × 100 mm, 1.8 μm) equipped with a Vanguard pre-column and maintained at 40 ºC. The column was eluted with (A) 0.1% formic acid in water and (B) 0.1% formic acid in acetonitrile at a flow rate of 0.5 mL min^-1^. The gradient was programmed as follows: start 1% B for 1 min, ramp to 15% B from 1-3 min, ramp to 50% from 3-6 min, ramp to 95% B from 6-9 min, hold until 10 min, ramp to 1% from 10-10.2 min, and a final hold at 1% B (total gradient time 12 min). Separate autosampler injections of 5 μL each were made for positive and negative ion modes.

The samples were analyzed in a random order with a pooled sample run after every six samples. The mass spectrometer was operated in selected reaction monitoring (SRM) mode; optimal SRM parameters (s-lens, collision energy) for each target compound were optimized individually using an authentic standard^37^. Two SRM transitions per compound were monitored for quantification and confirmation. Eight-point external calibration curves based on peak area were generated for each compound. The resulting data were converted to mzML files using the msConvert tool^38^and processed with MAVEN^39^.

## Acknowledgements

We thank Gabriel E. Leventhal, Shaul Pollak Pasternak and Elizabeth Kujawinski for their comments and careful reading of the manuscript. We also thank all members of the Cordero lab, and in particular José Saavedra and Matthew Metzger for their support. This project was supported by Simons Early Career Award 410104, the Alfred P Sloan fellowship FG-20166236 and the Simons Collaboration: Principles of Microbial Ecosystems (PriME), award number 542395.

## Contributions

MSD, NC and OXC designed the study. MSD, NC, TNE, JS, DS, and JB executed the study. MSD, TNE, JS, DS and OXC analyzed the results. TNE and OXC wrote the paper.

